# 3D organoid modeling identified that targeting IGF1R signaling may overcome drug resistance in breast cancer

**DOI:** 10.1101/2023.05.14.540701

**Authors:** Ekansh Mittal, David Qian

**Affiliations:** Oregon Health & Science University, Portland, OR USA, 97239

## Abstract

Breast cancer is the most frequently diagnosed cancer and the second largest cause of cancer deaths in women. However, drug resistance and poor response to treatments are common. Thus, there is an unmet need to identify new treatments and effective lab-based drug testing methods. Here we established a novel 3-dimensional organoid method by co-culturing cancer cells with healthy endothelial cells for longer-term testing of new drug combinations that combat drug resistance. As a proof-of-concept we showed that paclitaxel efficacy can be improved by combining it with AKT inhibitors. In addition, we identified a new triple combination of paclitaxel, HER2 inhibitor, and IGF1R inhibitor, which is more effective in increasing cell death and reducing organoid growth. Interestingly, many IGF1R pathway members are upregulated in breast cancer patients, and high expression is associated with poor survival, indicating that IGF1R is an attractive therapeutic target. Overall, using this novel organoid method, we can mimic more accurate culture conditions and identify new targets to be tested in clinical trials. Our approach is applicable to various cancers to improve patients’ outcomes.

## Introduction

Breast cancer is the most frequently diagnosed cancer and the second cause of cancer deaths in women. In 2022, 287,850 women were estimated to be diagnosed with invasive breast cancer in the United States and approximately 43,250 women were estimated to have died of their disease^1^. Men can also have breast cancer; however, male breast cancer is rare. Despite of all the efforts, these estimated numbers are significantly higher than the cases diagnosed in prior years^2-5^, demanding improvement in existing methods for therapeutic target identification. The breast cancer has four molecular subtypes: (1) luminal A, (2) luminal B, (3) human epidermal growth factor receptor 2 (HER2)-enriched, and (4) Triple-negative (basal-like), which have critical differences in incidence, response to treatment, disease progression, survival, and imaging features. Luminal tumors are most common, characterized by estrogen receptor (ER) expression. Luminal A has the best prognosis of all subtypes, whereas patients with luminal B tumors have significantly shorter overall and disease-free survival^6^. Thus, the identification of novel targets to overcome drug resistance is critical.

Further, it takes about 10 to 15 years and about $1 billion to develop one successful drug and despite all these enormous investments in time and money, over 90% of the drugs fail to meet objectives in clinical trials^7^. Some of these drugs fail because they didn’t treat the conditions they aimed to treat or the side effects were too strong. Consequently, many drugs never reach the approved state and thus wasting all the investments in time, money, and human resources. Most of the in vitro drug screening approaches use culturing of cancer cells in plastic dishes, which do not represent three-dimensional (3D) microenvironment and long-term culture effects. In vivo, models are informative but expensive and time taking, does not allow multiple screening at the same time, or may not represent human biology. Therefore, a proper drug screening strategy might help to improve the identification of effective targets. Here we established a 3D organoid model comprising cancer and healthy endothelial cells to test the efficacy of various inhibitors and identified that IGF1R targeting in combination with paclitaxel and HER2 might be more effective in inhibition the growth of cancer cells.

## Materials and methods

### Cell lines

MDA-MB-231 cell line (Triple-negative Breast Cancer, HTB-26) and HUVEC (Human Umbilical Vein Endothelial Cells, CRL-1730) were obtained from Oregon Health & Science University and were originally purchased from American Type Culture Collection (ATCC). MDA-MB-231 cell lines were cultured in DMEM media supplemented with 10% fetal bovine serum (FBS), 2 mM L-glutamine, and 100 U/ml penicillin, and 100 µg/mL streptomycin at 37ºC and 5% CO_2_. HUVEC cells were maintained in growth factor-supplemented media (EBM + EGM-2 aliquots + 10% FBS) from Lonza (C-3121, CC-4542). MDA-MB-231 cell line was modified to express GFP (green fluorescent protein) and HUVEC cells were modified to express RFP (red fluorescent protein).

### Drug Dilutions and MTS cell viability assay

The small molecule inhibitors such as paclitaxel (chemotherapy drug), IGFR1 inhibitor (GSK1838705A), and AKT inhibitors (MK2206) were obtained from Laboratory stocks or purchased commercially from MedChemExpress. The highest concentration of the drug 10 μM and the lowest being 13.7 nM for 7 points dose curve and included DMSO-only control for the vehicle arm. In the case of drug combinations, equal concentrations of two drugs were mixed during dilutions, which means 5 μM of paclitaxel with 5 μM of AKT or IGF1R inhibitors. The cells were plated at a density of 10,000 cells per well in the presence of a 7-dose concentration series with a total volume of 100 μl and cultured for 72 hours as described previously^8^. Next, cells were incubated with 20 μl of 4,5-dimethylthiazol-2-yl-2,5-diphenyltetrazolium bromide (MTT) (AqueousOne, Promega) at 37°C for 1-2 hours for MTS colorimetric cell viability assay. Blank wells with media and MTS (but no cells prepared) to calculate the background coming from MTS alone. The plates were read at an optical density of 490 nm to measure viability using a BioTek microplate reader instrument.

### Three-Dimensional (3D) Organoid culture

For organoid cultures, a modified protocol was used from the existing ones^9^. The breast cancer cells and HUVEC cells 50,000 each were mixed in 1:1 ratio in 100uL/well media along with Collagen I: 8 μg/mL (Stem Cell) and 1% Geltrex. 100k cells/well were plated in 96 well low-adhesion round bottom plates (Corning® 96-well Black/Clear Round Bottom Ultra-Low Attachment Spheroid Microplate).

### Organoid analysis for apoptosis by flow cytometry

The single-cell suspension of organoids prepared by digesting organoids using accutase at 37 °C for 5 mins. For flow cytometry-based analysis for apoptosis assay, single-cell suspension organoids were transferred to a deep-96 well plate that holds two ml volume and stained with CD44, CD24, and CD31 surface antibodies, along with annexin V and 7ADD. CD44 and CD24 are present on breast cancer cells^9,10^ and CD31 is present on endothelial cells^11,12^, which allowed the gating of both cell types. Cells were stained with surface antibodies in PBS with 1% BSA and incubated in the dark for 20 mins. Cells were washed twice at 1,500 RPM and resuspended in 100 μl Annexin V Binding Buffer and stained with 7ADD and Annexin V following manufacturer guidelines. The data was collected by flow cytometry on the Foretessa machine. Data was analyzed using Flow jo analysis.

### Organoid analysis by imaging

For image analysis of organoids, the culture media was gently removed, and washed once with PBS. The staining was performed using Hoechst 33324. The images were collected using Zeiss spinning disk confocal microscope for red, green, and blue fluorescent channels with z-stacking at 4 hr time lipase as well as statistic images. The area of the organoid was calculated using Zen software and intensity overtime was calculated using Imaris software.

### TCGA analysis

C-Bibportal^13^ was used to perform survival analysis for breast cancer patients with altered IGF1R signaling pathway and to analyze mRNA gene expression data for IGF1R signaling molecules.

### Statistical Analysis

p-value < 0.05 and FDR (false discovery rate) were used to determine the significance. The GraphPad Prizm and Excel were used for graphing and calculating mean, standard deviation, and p values.

## Results

### Establishing a novel three-dimensional (3D) organoid model for cancer cells for testing drugs

The above cell viability assay showed improvement in drug sensitivity but also showed that paclitaxel alone is very effective using this approach. Previous studies suggest that culturing cells in plastic dishes may not represent accurate culture conditions and require testing in a model that is three-dimensional. Therefore, we established a novel 3D in vitro culture method by co-culturing breast cancer (MDA-MB-231, Triple-negative Breast Cancer cells) and endothelial (HUVEC) cells. We were able to see the 3D organoid formation within 2-3 days and were able to culture organoids for 2-3 weeks **(Figure 1)**. Next, we used these organoids cultures for testing the efficacy of standard-or care treatment paclitaxel. We also tested the efficacy of paclitaxel in combination with AKT inhibitors which are being recently tested in clinical trials.^14^ We determined the IC50 for these drugs using cell viability assay to determine the concentrations to use in organoid models. Our results show that combining AKT inhibitor with paclitaxel reduced IC50 from 20 nM to 6 nM, respectively, and the area under the curve (AUC) from 230 nM to 183 nM, respectively. Based on these results in our initial organoid cultures, we used IC50 values for paclitaxel (20 nM) and AKT inhibitor (3.5 μM) **(Figure 2A)**. These concentrations had no effect on cell viability in organoid culture assay even after one week culturing, indicating culturing on 2D versus 3D may have variability in sensitivity. Therefore, we increased concentration for paclitaxel (80 nM) and AKT inhibitor (5.0 μM) for subsequent experiments. Our preliminary data using flow cytometry analysis showed that combinations of AKT inhibitor with paclitaxel lead to a significant increase in cell death (> 20% increase) as measured by apoptosis of CD44^+^ or CD44^+^CD24^+^ double-positive cells as compared to one or both single treatments. Interestingly these treatments have only a slight or no impact on CD31+ endothelial cells representing healthy cells from the microenvironment **(Figure 2B)**. Similarly, imaging analysis showed that the combined treatment with AKT inhibitor and paclitaxel led to a significant reduction in the growth of organoids size after 7 days of treatment compared to single treatments approximately 40-50% **(Figure 2C)**. These results show the feasibility of using organoid model as functional assays for testing drug combinations.

**Figure 1:**
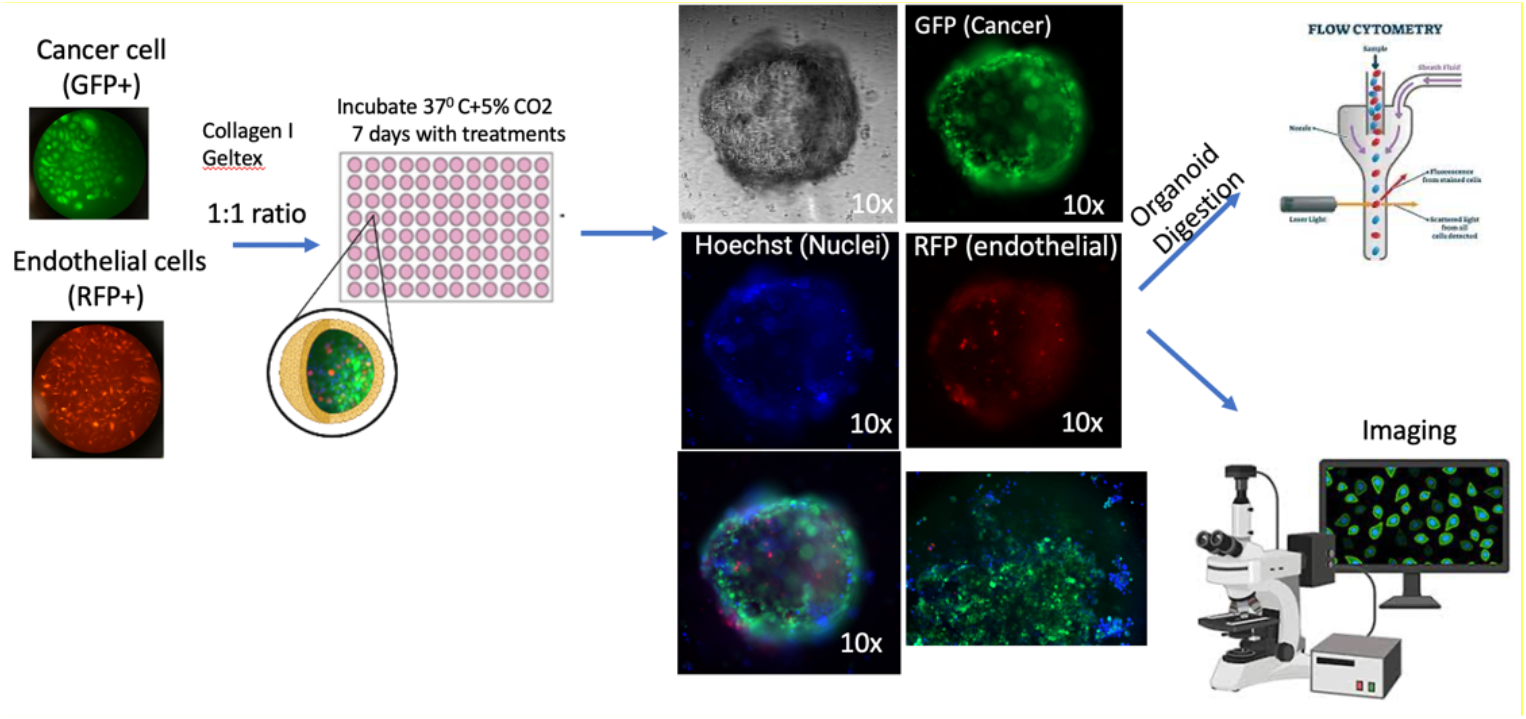
3D Organoid culture method: The breast cancer cells and HUVEC cells 50,000 each were mixed in 1:1 ratio in a 96 well low-adhesion round bottom plates Organoids were treated and fed twice per week. Organoids were analyzed by flow cytometry and by imaging analysis.

**Figure 2.**
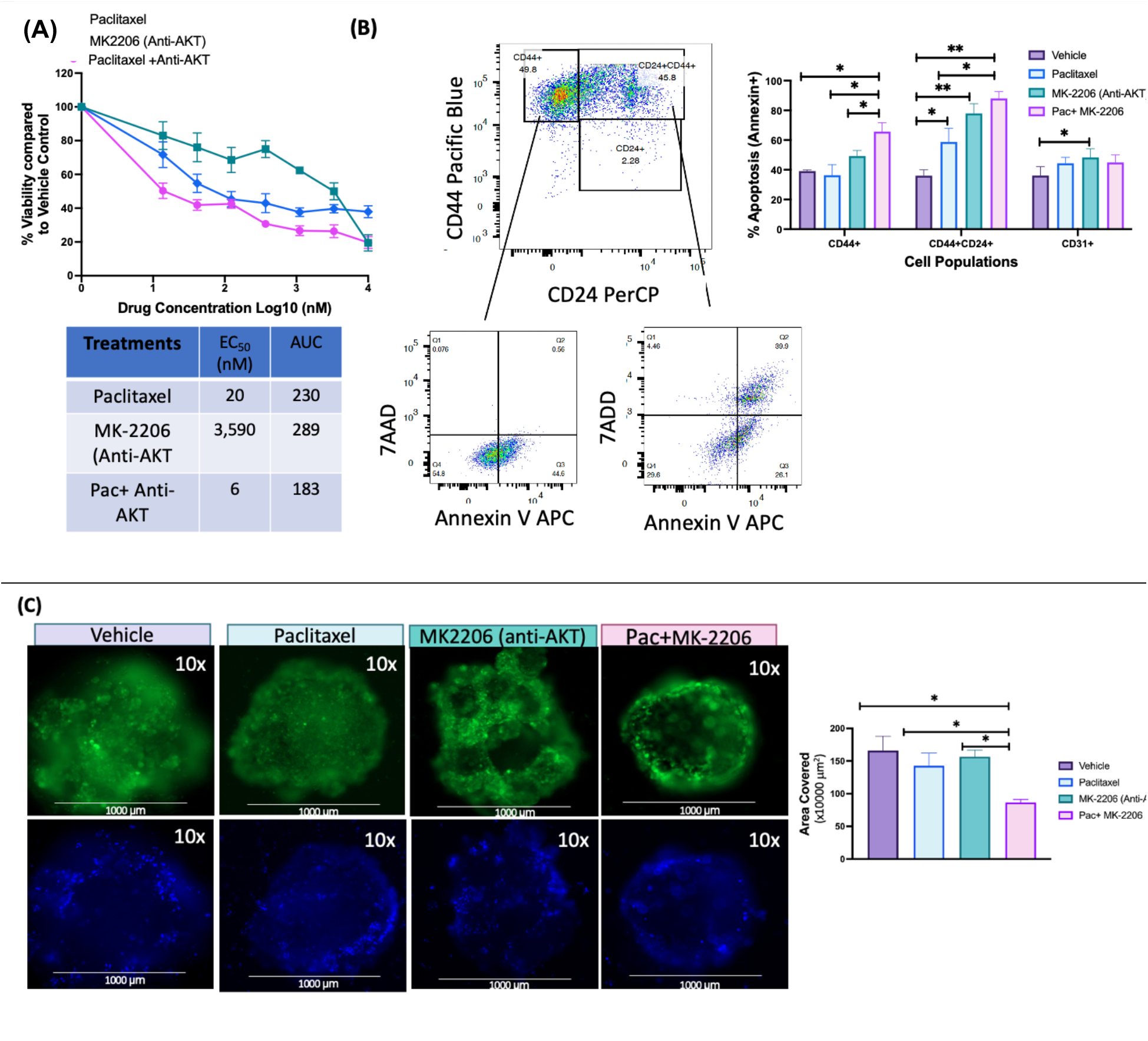
3D Organoid cultures utilized for functional screening of drug combinations. (A) Serial drug dilutions were established as described in methods using 7-point curve with maximum drug concentrations (10,000 nM equivalent to Log_10_ = 4 nM) and cells were cultured for 72 hrs and analyzed by MTS cell viability assay. **(B**) The organoids were digested and analyzed for apoptosis of CD44, CD44/CD24 double positive, and CD31+ cells using Flow cytometry methods. The data is represented as bar graph from triplicate experiments. **(C)** 10x images were analyzed using Zen software for total surface area of organoids and the experiment was performed twice in duplicates. P value represents * 0.05, ** 0.01.

### Testing the triple combinations using organoid cultures

TCGA differential expression analysis suggest that IGF1R signaling is critical for both basal and HER2-positive breast cancer. Further, the patients with upregulated IGF1 signaling have poor overall survival **(Figure 3)**.

**Figure 3.**
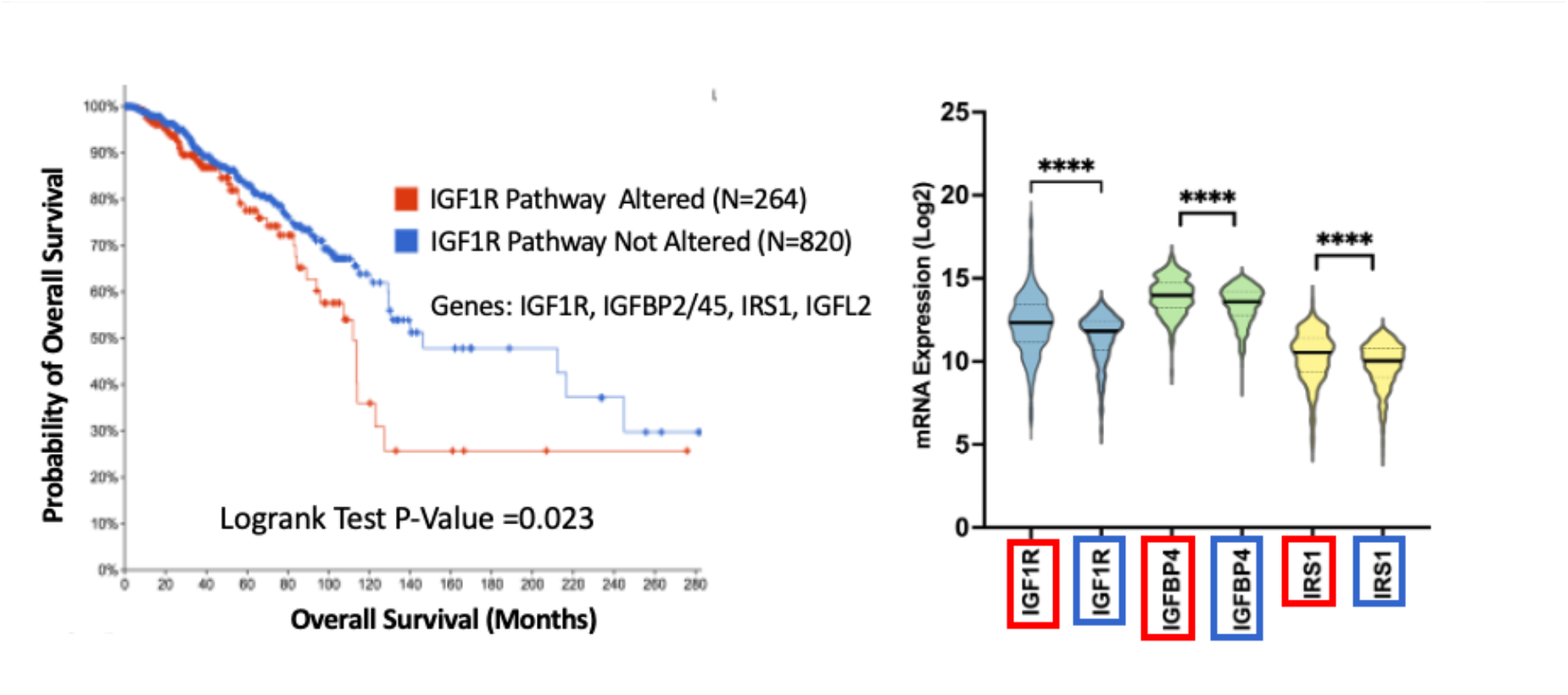
TCGA data analysis showing high expression of IGF1R signaling is correlated with poor overall survival of breast cancer patients (Left Panel). The expression of IGF1R signaling molecule (Right panel). An unpaired T-test was used to compare expressions of altered and unaltered arms.

Therefore, we further focused our work in testing the combination of IGF1R inhibitor with and without chemotherapy and HER2 inhibitor as double and triple combinations. The testing of IGF1R inhibitor in combination with paclitaxel and HER2 inhibitor showed lower area under curve (AUC) compared to each treatment alone when we tested these combinations (Figure 4A). 3D organoid analysis by flow cytometry showed increased apoptosis using in combination with single and double drug combinations of paclitaxel or HER2 or IGF1R inhibitors for CD44+ or CD24+CD44+ cancer cells, while no significant cell death was observed using HUVEC CD31+ cells (Figure 4B). Finally live imaging analysis overtime at week 1 and week 2 treatments show significant reduction in Hoesch positive cells in triple combination with paclitaxel+anti-HER2+anti-IGF1R showed the lower cells survival over two weeks. Interestingly, HER2 inhibitor itself is not effective as expected but in combination with paclitaxel showed growth reduction at week one, but that effect was not persistent over two weeks, while the combination of paclitaxel+anti-HER2+anti-IGF1R effect on growth remain persistent overtime. These results suggest that 3D organoid models can help identify effective drug combinations with persistent effects.

**Figure 4.**
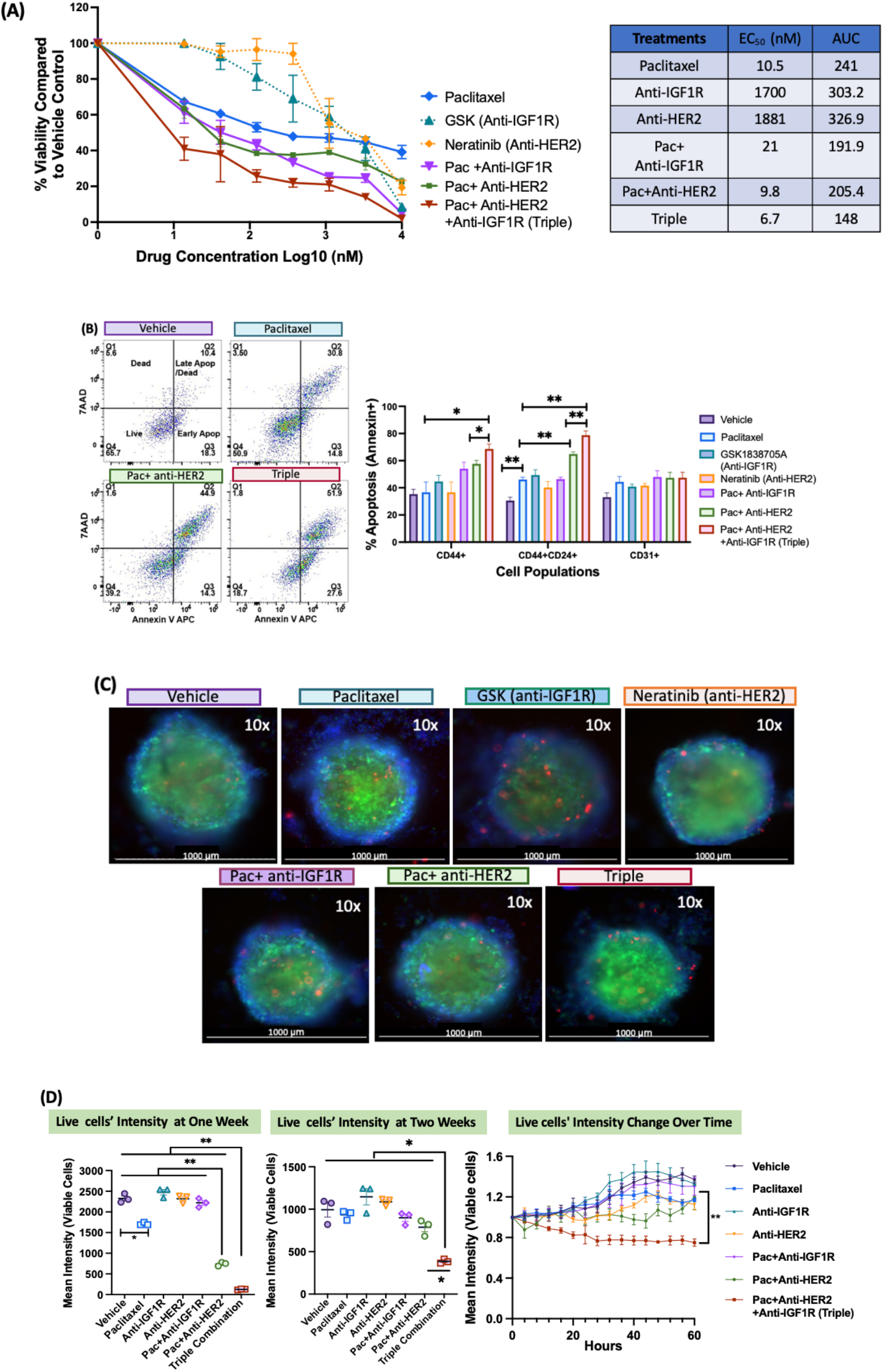
IGF1R inhibition in combination with chemotherapy and HER2 inhibitor is more sensitive compared to the single agents. (A) Drug dependency for a breast cancer cell line is shown for indicated drugs as single and combination treatments. Serial drug dilutions were established using 7-point curve with maximum drug concentrations and cells were cultured for 72 hrs and analyzed by MTS cell viability assay. The data is represented as percentage viability. EC50 and Area Under Curve (AUC) was calculated using Graphpad. **(B)** The organoids were cultured for two weeks in paclitaxel (80 nM) or HER2 inhibitor (Neratinib. 2.0 μM) or IGF1R inhibitor (GSK1838705A 2.0 μM) or combinations for two weeks. Organoids were dissociated and analyzed for apoptosis of CD44+, CD44/CD24+ and CD31+ cells using Flow cytometry. The data is represented as a bar graph from triplicate experiments. **(C)** 10x Images of individual Organoids. (D) The live cell imaging of 3D organoids was performed for 60 hrs at week 1 and week 2 using Spinning Disc Confocal Microscope with 4 hrs time-lapse and 46 Z-slices. Images were processed using Imaris Software. The mean intensity of live cells (Hoesch staining, blue Channel) is shown at week one by fold change over 60 hrs (right panel), absolute mean intensities are shown at weeks one and two (middle and right panels). The experiment was performed twice (Total three replicates). p values * < 0.05, ** <0.01 using multiple paired T test.

## Discussion

Cancer is a leading cause of death worldwide and the most important impediment to increasing life expectancy in every country of the world in the 21st century^1^. Recently there has been a small decrease in death rates in colorectal, prostate, and female breast cancer largely due to advances in early detection and more effective treatments^1^. Further, the success rate for the oncological clinical trials is only 3.4%. The lack of failure of these trials is limited efficacy and increased toxicity, suggesting that more precise approaches are needed for new target identifications and to improve patient outcomes.^15^ Therefore, we developed a 3D cell culture method that can mimic cancer microenvironment more closely. We used this model to show the efficacy of standard-of care drugs and to identify new effective combinations.

Initially, we used commonly used cell viability-based colorimetric assay, where cells are grown on plastic plates over time. This assay showed the high efficacy of paclitaxel (standard of care treatment) in breast cancer cell lines and the same is supported by the database search for 50 cell lines that paclitaxel has efficacy in low nanomolar in cell line models using this assay^16,17^. However, we know from clinical trials and treating patients that drug resistance is very common to paclitaxel and alone might not be that effective. Literature also suggests that the microenvironment plays a critical role in drug resistance and culturing cells just on plastic in 2-dimension might not accurately represent what we observe in vivo. Therefore, we thought of establishing the 3-dimension organoid culture method. Similar methods were previously used to develop organoid methods in vivo using mouse models for breast cancer. But in vivo testing for these many targets and many concentrations are not feasible to test simultaneously. Therefore, I thought to establish breast cancer organoid models in vitro using a 96-well format. Using the organoid culture method, I was able to test one combination that is in a clinical trial (paclitaxel +MK2206) and a new combination of paclitaxel with IGF1R inhibitor and with and without HER2 inhibitor. Both combination testing in organoid model shows that single treatments are not that effective compared to combined treatments, thus mimicking actual breast cancer patients more closely. Use of HUVEC endothelial cells in culture allowed us to test simultaneously if treatment could be toxic to healthy cells.

Upon further validation, these targets can lead to new treatment options for aggressive breast cancer subtypes such as Triple-negative breast cancer. In the future, this approach will potentially be applied to any cancer type. Overall, this approach can result in more effective and accurate decision-making, reduce the time of testing each drug and target in the laboratory setting, and possibly narrow down the target list that could be most effective. This could save a significant amount of time and money and provide a better cure for the patients, thus can improve their quality of life and disease outcomes.

## Authors Contributions

EM has put together the concept, collected the data, and performed the analysis, prepared first draft the manuscript. DQ provided guidance throughout and gave feedback for the project and manuscript.

## Acknowledgments

OHSU imaging and flow cytometry core for their support, and lab members for providing supervision and guidance for methods learning.

